# The Temporal Pattern of Spiking Activity of a Thalamic Neuron are Related to the Amplitude of the Cortical Local Field Potential

**DOI:** 10.1101/2021.09.08.459532

**Authors:** Hiroshi Tamura

## Abstract

Neuron activity in the sensory cortices mainly depends on feedforward thalamic inputs. High-frequency activity of a thalamic input can be temporally integrated by a neuron in the sensory cortex and is likely to induce larger depolarization. However, feedforward inhibition (FFI) and depression of excitatory synaptic transmission in thalamocortical pathways attenuate depolarization induced by the latter part of high-frequency spiking activity and the temporal summation may not be effective. The spiking activity of a thalamic neuron in a specific temporal pattern may circumvent FFI and depression of excitatory synapses. The present study determined the relationship between the temporal pattern of spiking activity of a single thalamic neuron and the degree of cortical activation as well as that between the firing rate of spiking activity of a single thalamic neuron and the degree of cortical activation. Spiking activity of a thalamic neuron was recorded extracellularly from the lateral geniculate nucleus (LGN) in male Long-Evans rats. Degree of cortical activation was assessed by simultaneous recording of local field potential (LFP) from the visual cortex. A specific temporal pattern appearing in three consecutive spikes of an LGN neuron induced larger cortical LFP modulation than high-frequency spiking activity during a short period. These findings indicate that spiking activity of thalamic inputs is integrated by a synaptic mechanism sensitive to an input temporal pattern.

**Significance Statement:** Sensory cortical activity depends on thalamic inputs. Despite the importance of thalamocortical transmission, how spiking activity of thalamic inputs is integrated by cortical neurons remains unclear. Feedforward inhibition and synaptic depression of excitatory transmission may not allow simple temporal summation of membrane potential induced by consecutive spiking activity of a thalamic neuron. A specific temporal pattern appearing in three consecutive spikes of a thalamic neuron induced larger cortical local field potential modulation than high-frequency spiking activity during a short period. The findings indicate the importance of the temporal pattern of spiking activity of a single thalamic neuron on cortical activation.

## Introduction

The activity of neurons in sensory cortices mainly depends on feedforward thalamic inputs (Ferster et al., 1996; Chung and Ferster, 1998; Liu et al, 2007; Li et al., 2013; Lien and Scanziani, 2013). Because a single thalamic spike induces a small depolarization in the postsynaptic cortical neuron (Gil et al., 1999; Stratford et al., 1996; Brecht and Sakmann, 2002; Bruno and Sakmann, 2006; Lien and Scanziani, 2018; Sedigh-Sarvestani et al., 2019; Ringach, 2021), synaptic inputs should be summated to allow the membrane potential to reach the spiking threshold. Synaptic inputs can be summated temporally (Magee, 2000; Feldmeyer et al., 2002). If this is the case in thalamocortical synapses, high-frequency spiking activity during the short period of a thalamic neuron is associated with larger cortical activation than that of a single spike (Usrey et al., 2000).

However, the presence of feedforward inhibition (FFI) at thalamocortical synapses (Ferster and Lindström, 1983; Gil and Amitai, 1996; Swadlow, 2002; Beierlein et al., 2003; Gabernet et al., 2005; Kimura et al., 2010; Ji et al., 2016; Bereshpolova et al., 2020) may not allow effective temporal summation of thalamic inputs by a cortical neuron. Thalamocortical relay neurons directly synapse onto inhibitory cortical neurons that inhibit cortical layer 4 neurons (Gil and Amitai, 1996; Beierlein et al., 2003; Gabernet et al., 2005; Kimura et al., 2010). Thus, FFI creates a short temporal window for synaptic integration (Gabernet et al., 2005; Kimura et al., 2010) and suppresses depolarization induced by the latter part of high-frequency spiking activity from thalamocortical relay neurons. Furthermore, the efficacy of thalamocortical excitatory synapses is depressed during repetitive activation of thalamic inputs, with subsequent spikes of a thalamic neuron inducing much smaller depolarization than the first spike (Gil et al., 1997; Feldmeyer et al., 2002; Beierlein et al., 2003). Therefore, the high-frequency spiking activity of a thalamic neuron is not as effective as that expected from the simple temporal summation of excitatory synaptic inputs.

The spiking activity of a thalamic neuron in a specific temporal pattern may circumvent the FFI and depression of thalamocortical excitatory synapses. For example, during a long period of silence of a thalamic neuron, the inhibitory current induced by FFI was decreased and the depression of excitatory synapses was alleviated. Indeed, a single spike after a long period of silence of a thalamic neuron was associated with large cortical activation (Swadlow and Gusev, 2001; Swadlow, 2002; Swadlow et al., 2002; Stoelzel et al., 2008; Stoelzel et al., 2009), suggesting specific temporal patterns of a single thalamic neuron can be used to ensure thalamocortical transmission.

The present study examined whether the firing rate or temporal pattern of spiking activity of a single thalamic neuron was related to the degree of cortical activation. The spiking activity of a thalamic neuron was extracellularly recorded from the lateral geniculate nucleus (LGN) with simultaneous recording of local field potential (LFP) from the visual cortex (VC) in male Long-Evans rats. LFPs represent neural activity from a cortical region of a few millimeters (Mitzdorf, 1987; Logothetis, 2003; Kreiman et al., 2006; Jin et al., 2008; Nauhaus et al., 2009; Kajikawa and Schroeder, 2011; Buzsáki et al., 2012; Pesaran et al., 2018), and are linearly related to the membrane potential and firing rate of cortical neurons (Deweese and Zador, 2004; Poulet and Petersen, 2008; Okun et al., 2010; Lien and Scanziani, 2013). To examine the relationship between the spiking activity of an LGN neuron and cortical LFP, the LGN spike-triggered average (STA) of LFP was calculated. Although both firing rate and temporal pattern of spiking activity of an LGN neuron were related to amplitude of STA-LFP recorded from VC, a specific temporal pattern of spiking activity appearing in three consecutive spikes of an LGN neuron induced larger LFP modulation recorded from VC than high-frequency spiking activity during the short period. The importance of temporal pattern was confirmed with the analysis of monosynaptically connected LGN-VC neuron pairs.

## Materials and Methods

The general experimental procedures were previously described (Kimura et al., 2010; Ikezoe et al., 2012). Neuronal activity was recorded from 17 adult male Long-Evans rats (260–360 g bodyweight). All experiments were performed in accordance with the guidelines of the National Institute of Health (1996) and the Japan Neuroscience Society, and were approved by the Osaka University Animal Experiment Committee (FBS-18-003).

### Neuronal activity recording

Rats were anesthetized using urethane (Sigma-Aldrich, Tokyo, Japan; 1.2–1.4 g/kg, intra-peritoneal injection) with urethane supplementation if necessary. The head was restrained with ear bars coated with local anesthetic (2% lidocaine; AstraZeneca, Osaka, Japan). Local anesthetic (0.5% lidocaine; Maruishi, Osaka, Japan) was administered into the scalp before incision. A small hole (approximately 4 mm in diameter) was drilled in an appropriate part of the skull and a small slit (1–3 mm) was made in the dura to insert an electrode. Thalamic spiking activity was recorded from neurons in the LGN using a single-shaft multiprobe electrode (A1×32-Poly3-10mm-50-177; NeuroNexus, Ann Arbor, MI, USA). The electrode was equipped with 32 recording probes arranged in three columns at the tip; the center column had 12 probes and the left and right columns had 10 probes each. The distance between the centers of the adjacent recording probes was 50 μm. The LGN electrode was inserted into the brain at 4.5 mm posterior to the bregma and 5.6 mm lateral from the midline (Fig. 1*A*). The penetration angle was 30° in the coronal plane. A reference electrode was positioned on the surface of the scalp. LGN recordings were made at average intervals of 0.21 mm along a penetration and were typically performed three times per penetration.

**Figure 1.**
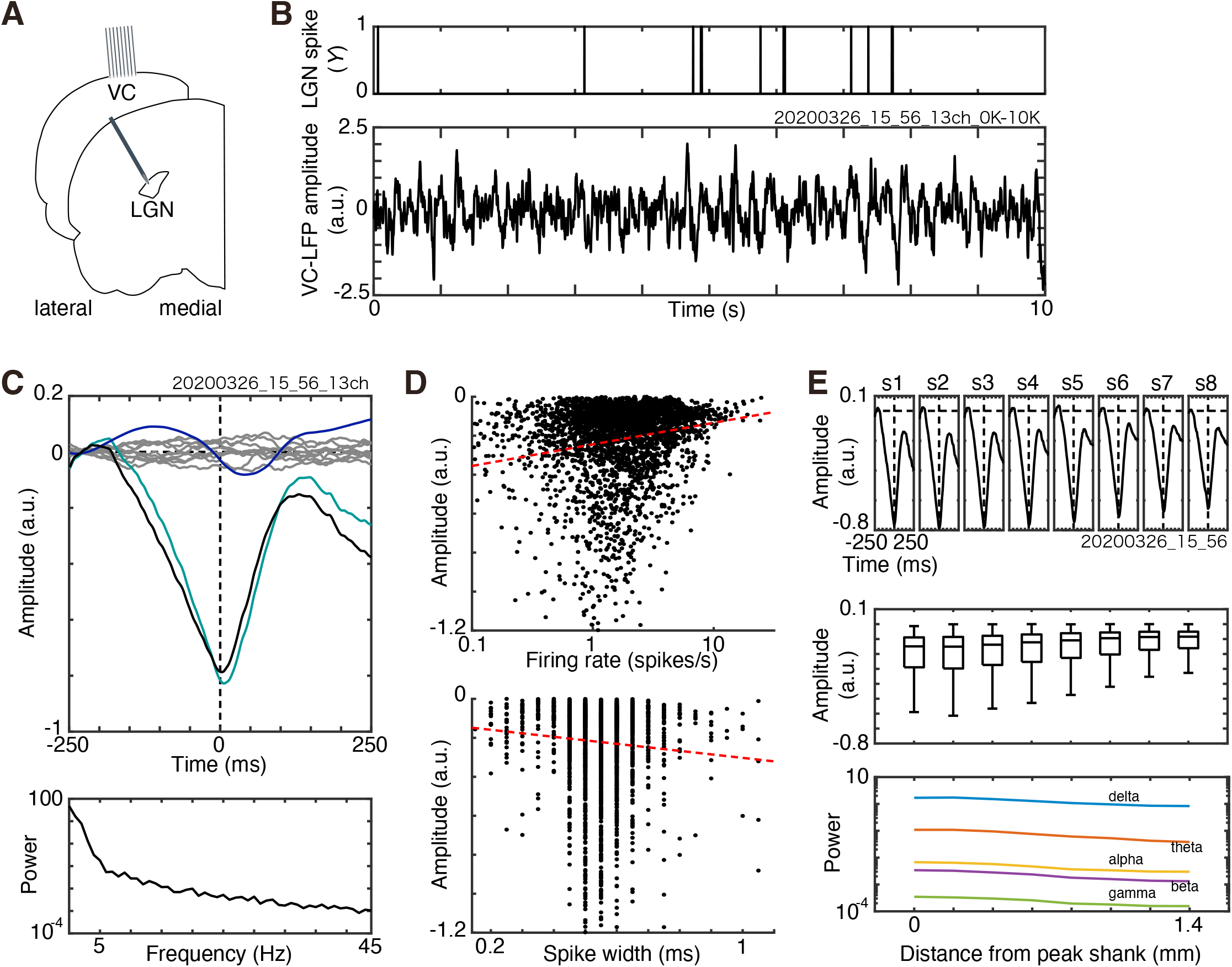
The lateral geniculate nucleus (LGN) neuron spike-triggered average (STA) of the cortical local field potential (LFP). ***A***, Schematic representation of the recording configuration. The electrodes for LGN and the visual cortex recordings are depicted on coronal sections of a rat brain. Note that the brain sections and the two electrodes are not drawn to the exact scale for illustrative purposes. ***B***, An example of a spike train of an LGN unit (top) and simultaneously recorded cortical LFP (bottom). The vertical lines in the top panel show the timing of the LGN spikes. If a spike occurs *Y* = 1, otherwise *Y* = 0. The presented cortical LFP was spike-removed, low-pass filtered, down-sampled, and z-score transformed. ***C***, Top: an example of STA-LFP. The green, blue, and black lines represent the raw STA-LFP, peristimulus time histogram-predicted STA-LFP, and subtracted STA-LFP, respectively. The ten gray lines represent raw STA-LFPs calculated with ten randomized spike trains. Average amplitudes from − 250 to − 200 ms were subtracted to align the signals. The vertical dashed line at 0 ms represents the timing of the triggering-LGN spike. The horizontal dashed line indicates zero amplitude. Bottom: power spectra of the subtracted STA-LFP in the top panel. ***D***, Top: relationship between the average firing rate during the total recording period of an LGN unit and the STA-LFP amplitude. Each dot represents a single LGN unit. The red dashed line represents a linear regression. Bottom: relationship between the spike width of an LGN unit and the STA-LFP amplitude. ***E***, Top: an example of STA-LFPs recorded from the eight shanks (s1–s8) of a cortical electrode. Middle: relationships between the horizontal distance from the shank (peak shank) that recorded the STA-LFP with the maximum amplitude *and* STA-LFP amplitude. In the box plot, the center of each box (black horizontal lines) represents the median across LGN units, whereas the top and bottom of the box represent the upper and lower quartiles, respectively. The attached whiskers connect the most extreme values within 150% of the interquartile range from the end of each box. Bottom: relationships between the horizontal distance and power of each frequency component of STA-LFP (blue, *δ* (1–3 Hz); orange, *θ* (4–7 Hz); yellow, *α* (8–11 Hz); purple, *β* (12–29 Hz); green, *γ* (30–250 Hz)). The median across LGN units was plotted.

The spiking activity and LFP from the VC was recorded using an eight-shank electrode (Buzsaki64; NeuroNexus). The distance between the centers of adjacent shanks was 200 μm. Each shank was equipped with eight recording probes arranged in a V-shape at the tip. The distance between the centers of adjacent recording probes was 20–40 μm. The center of the electrode for recording from the VC was located at 6.5 mm posterior to the bregma and 3 mm lateral from the midline (Fig. 1*A*). The electrode plane was parallel to the coronal plane. The penetration angle was adjusted so that the tips were parallel to the cortical surface as much as possible. A reference electrode was positioned on the surface of the scalp. Once the spiking activity was obtained from most of the shank, insertion of the cortical electrode was stopped and the position was maintained during recording.

Recordings from the LGN and VC were performed in the right hemisphere. Signals were amplified (10,000×), filtered (1.3–7,603.8 Hz), digitized (20,000 Hz; RHD 2000 amplifier board; Intan Technologies, Los Angeles, CA, USA), and stored in a personal computer for offline analysis.

Visual stimuli were presented to enhance spiking activity of LGN neurons and obtain a reliable STA-LFP from a limited recording duration. Visual stimuli were presented to the left eye (i.e., contralateral to the recording hemisphere) in a bright room on a liquid crystal display (XL2546-B ZOWIE; BenQ, Taipei, Taiwan) placed 30 cm from the animal’s left eye at 30° from the anterior-posterior body axis. The receptive field (RF) positions of the LGN and VC recording sites were qualitatively determined by monitoring the spiking activity of multiunits while manually presenting grating stimuli. Visual stimuli were presented to include the manually determined RFs. The stimulus set consisted of 200 one-dimensional noise square-wave gratings (spatial frequency, 0.016–0.15 cycles/°). Note that LGN and V1 neurons prefer a spatial frequency of 0.03–0.06 and 0.04–0.15 cycles/°, respectively (Girman et al., 1999; Sriram et al., 2016). The stimulus size was 108°×55° (horizontal × vertical). The gratings had one of four orientations (0°, 45°, 90°, or 135°) and were black and white. The luminance values of the black and white pixels were 1.0 cd/m^2^ and 319 cd/m^2^, respectively. Each stimulus was presented 20 times in a pseudo-random order for 0.5 s without an inter-stimulus interval. Thus, neural activity was recorded for approximately 30 min. The stimulus timing was recorded via a photodiode attached to the display. Saline was applied to the eyes before and after the recording of neural activity.

In 3 of 17 rats, neural activity was also recorded without visual stimuli (gray background, spontaneous activity) for approximately 30 min.

### Data analysis

Single unit activity was isolated offline using Kilosort (Pachitariu et al. 2016). Spike-sorting results were verified by calculating the auto-correlogram and the cross-correlogram (see Tamura et al., 2014 for details).

To perform STA-LFP, raw signals recorded from the VC were processed as follows. First, signals associated with the spiking activity of cortical neurons were removed from the raw signals. For this, the average spike waveform (3-ms duration) was calculated from the raw signals for each single unit recorded from the VC and the average spike waveform was subtracted from the raw signals. This process was repeated for all the isolated units recorded from the VC. Next, the spike-removed signals were low-pass filtered (<100 Hz) and down-sampled from 20,000 Hz to 1,000 Hz. Finally, the z-score standardized LFP was calculated by subtracting the average and dividing with the standard deviation of the processed LFP.

To obtain the STA-LFP, the cross-correlation between the spike train of an LGN unit and the z-score standardized LFP recorded from the VC was calculated at a temporal resolution of 1 ms with a temporal window of ±250 ms (raw STA-LFP). Because visual stimulation can modulate the spiking activity of an LGN unit and LFP recorded from the VC, the effect of visual stimulation contaminates raw STA-LFP. The effect of visual stimulation was estimated by calculating the cross-correlation between the peristimulus time histogram (PSTH) of spiking activity of an LGN unit and the stimulus-triggered cortical LFP (PSTH-predictor). By subtracting the PSTH-predictor from the raw STA-LFP, the subtracted STA-LFP was obtained where the effect of visual stimulation was removed. This method was similar to that used for the cross-correlation analysis of spike trains collected during visual stimulation (Perkel et al., 1967; Toyama et al., 1981; Tamura et al., 2004). In the Results, the subtracted STA-LFP is presented as the STA-LFP unless otherwise specified. STA-LFP was calculated for all 64 LFPs recorded using 64 probes. The amplitude of STA-LFP was quantified by measuring its initial negative deflection. Statistical significance of the amplitude of STA-LFP was evaluated by comparing it with that of STA-LFP triggered by a randomized spike train. The rationale was that if a relationship between LGN spike occurrence and LFP modulation was independent, the amplitude of STA-LFP obtained with an original spike train would be similar to that obtained with a randomized spike train. A randomized spike train was generated by randomizing the timing of spike occurrence. This process was repeated 10 times. The amplitude of the STA-LFP from the original spike train were statistically compared with the amplitude of STA-LFP from 10 randomized spike trains (*p* < 0.05, signed-rank test, one-tailed).

The cross-correlation between the spike train of an LGN unit and that of a VC unit was calculated at a temporal resolution of 1 ms with a temporal window of ±250 ms and obtained raw cross-correlogram (raw CCG). Because visual stimulation can modulate the spiking activity of an LGN unit and a VC unit, the effect of visual stimulation contaminates raw CCG. The effect of visual stimulation was estimated by calculating the cross-correlation between the PSTH of spiking activity of an LGN unit and that of a VC unit (PSTH-predicted CCG). By subtracting the PSTH-predicted CCG from the raw CCG, the subtracted CCG was obtained where the effect of visual stimulation was removed (Perkel et al., 1967; Toyama et al., 1981; Tamura et al., 2004). In the Results, the subtracted CCG is presented. Statistical significance of CCG peak was evaluated by comparing raw CCG with that of PSTH-predicted CCG (*P* < 0.0001, binomial test).

### Statistical analysis

All data were pooled for statistical analyses. The number of units used for each experiment is described in the Results. All statistical analyses were performed using statistical software (MATLAB; The MathWorks, Natick, MA, USA). The statistical threshold for *p-*values was set at 0.01 unless otherwise specified. Statistical outcomes (e.g., the chi-square [χ²] value) are provided. The effect size for non-parametric tests between two groups was estimated using the formula: 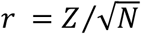, where Z and N represent the Z-statistic from the signed-rank test and the number of samples, respectively.

## Results

The present study was based on 4173 LGN units obtained from 47 recordings with visual stimulation and 713 LGN units obtained from 8 recordings without visual stimulation (spontaneous activity).

Spiking activity of an LGN unit induced clear modulation in LFP recorded from the VC. The spike train obtained from an LGN unit (Fig. 1*B*, top) was cross-correlated with the LFP recorded from the VC (Fig. 1*B*, bottom) to obtain raw STA-LFP (Fig. 1*C*, top, green line). By subtracting the PSTH-predictor (Fig. 1*C*, top, blue line) from the raw STA-LFP, the subtracted STA-LFP (Fig. 1*C*, top, black line) was obtained. The subtracted STA-LFP is presented as the STA-LFP unless otherwise specified. STA-LFP showed an initial negative deflection followed by a positive deflection. Most of the power of STA-LFP was in the low-frequency range (Fig. 1*C*, bottom).

The amplitude of STA-LFP was weakly related to the physiological properties of LGN neurons. Here, LFPs with the largest amplitudes in the STA-LFP among the 64 LFPs recorded from 64 probes were selected for each LGN unit and subjected to analysis. If the negative peak was not detected in STA-LFP, the unit was not included in the analysis. The average firing rate during the total recording period of an LGN unit was positively correlated with the amplitude of STA-LFP (Spearman’s correlation coefficient, *r* = 0.15, *p* = 3.70×10^− 19^, number of units = 3335; Fig. 1*D*, top). Because the negative deflection of STA-LFP was quantified, a positive correlation indicated an LGN unit with a lower average firing rate tended to induce larger modulation in STA-LFP. The spike width that is the duration measured from the timing of initial negative nadir to that of the subsequent positive peak of an LGN unit was negatively correlated with the amplitude of STA-LFP (Spearman’s correlation coefficient, *r* = − 0.10, *p* = 4.58×10^− 9^, number of units = 3335; Fig. 1*D*, bottom). These results suggest that LGN neurons with a lower average firing rate and broader spike width tended to induce larger modulation in STA-LFP.

The statistical significance of the amplitude of STA-LFP was examined by comparing the amplitude of STA-LFP triggered by the original spike train with that triggered by a random spike train. The amplitude of the STA-LFP in Figure 1*C* (top, black line) is significantly larger than that calculated with random spike trains (top, gray lines; *p* < 0.05, signed-rank test, one-tailed). Among 4173 LGN units obtained with visual stimulation, 3152 LGN units (76%) evoked significant negative modulation in STA-LFP. Among 713 LGN units obtained without visual stimulation, 500 units (70%) evoked significant negative modulation in STA-LFP.

The amplitude of STA-LFP decreased only slightly with horizontal distance. The analysis was performed using LGN units with significant STA-LFPs. The largest amplitudes among eight STA-LFPs recorded from eight probes on a shank represented the shank. An example of STA-LFPs recorded from eight shanks of the cortical electrode is shown in the top panel of Figure 1*E*. Clear modulation in STA-LFP was not limited to signals recorded from a single shank, but observed in those from all the eight shanks. The amplitude of STA-LFP recorded from a probe on the second shank (Fig. 1*E*, top, s2) was the largest, and that from the eighth shank (Fig. 1*E*, top, s8) was the smallest and was 83% of the maximum. The amplitude of STA-LFP decreased gradually from the peak with horizontal separation (Spearman’s correlation coefficient, *r* = − 0.73, *p* ≃ 0; Fig. 1*E*, middle), and the amplitude of STA-LFP recorded from the shank that was most distant from the peak shank was 67% of the maximum. The power of each frequency component (*δ*, 1–3 Hz; *θ*, 4–7 Hz; *α*, 8–11 Hz; *β*, 12–29 Hz; *γ*, 30–250 Hz) of STA-LFP had a similar tendency (Fig. 1*E*, bottom).

### The firing rate of LGN spikes is positively correlated with the amplitude of cortical LFP

The relationship between the firing rate of spiking activity of LGN neurons during a short period and the amplitude of cortical LFP was examined. STA-LFP was calculated for each firing-rate group, which was defined according to the number of spikes of a single LGN unit within a window of 20 ms (i.e., one-spike/20 ms, two-spikes/20 ms, three-spikes/20 ms, and four-spikes/20 ms; Fig. 2*A*). A 20-ms window was selected because it is close to the half-width of thalamocortical input-evoked excitatory postsynaptic potential (Gil and Amitai, 1996) and the membrane time constant of layer 4 regular spiking neurons (Beirlein et al., 2003; Gabernet et al., 2005). The first spike during the 20-ms period was used as the trigger spike for LFP averaging (Fig. 2*A*, thick vertical lines). LGN units that evoked significant modulation in STA-LFP were included in the analyses if at least 50 triggering spikes were recorded for each firing-rate group.

**Figure 2.**
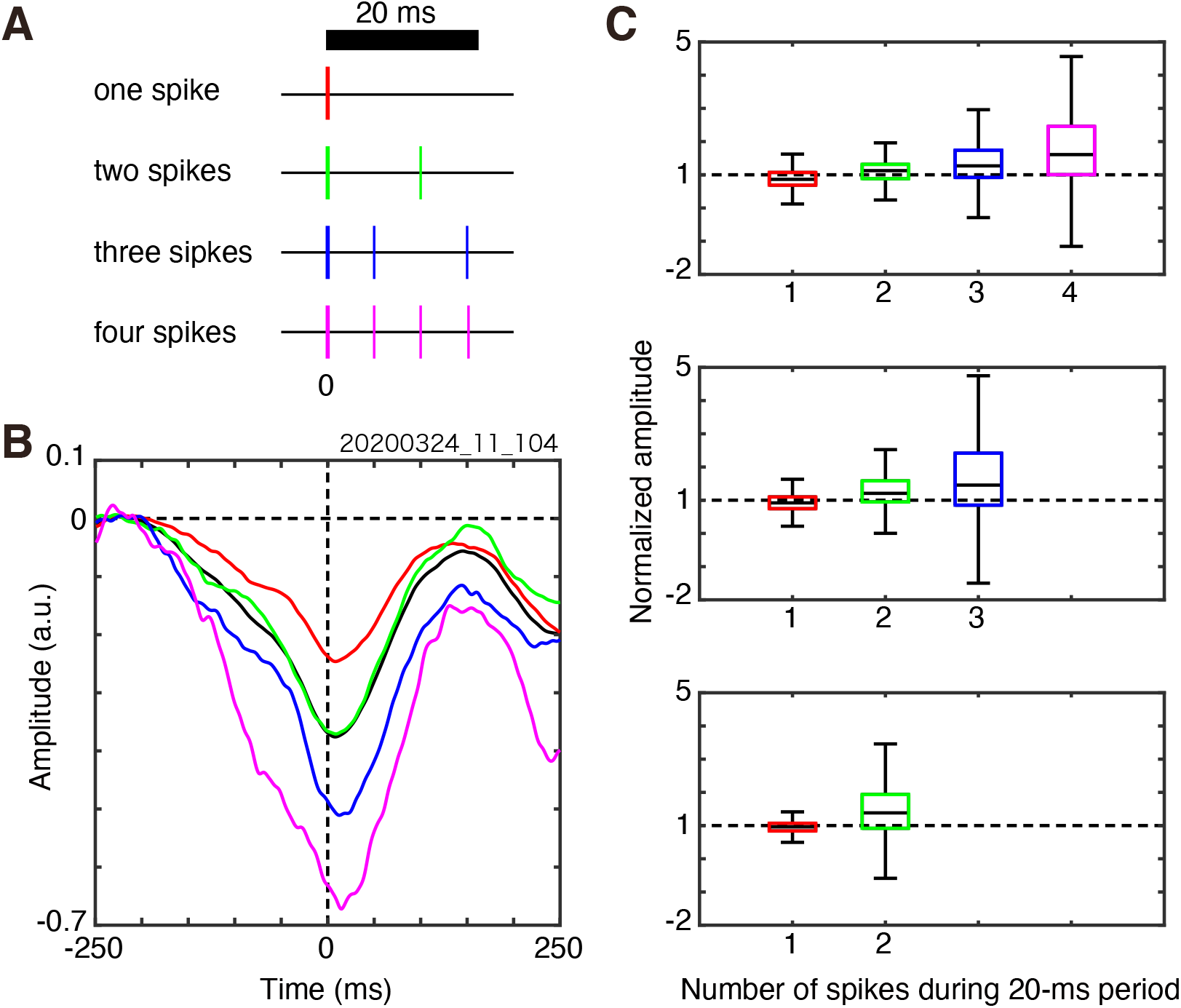
Relationship between the firing rates of a lateral geniculate nucleus (LGN) neuron during a 20-ms period and the cortical local field potential (LFP) amplitude. ***A***, Triggering spikes of an LGN unit were classified into one of four firing-rate groups according to the number of spikes within a 20-ms period (one spike, two spikes, three spikes, and four spikes). The triggering spike was the first spike within a 20-ms period (indicated by thick vertical lines at time zero). The black horizontal bar at the top represents the 20-ms period. ***B***, An example of spike-triggered average (STA)-LFPs calculated with an LGN unit for each firing-rate group (red, one spike; green, two spikes; blue, three spikes; magenta, four spikes). The STA-LFP calculated with all spikes of the LGN unit was also plotted (black line). ***C***, Comparisons of the normalized STA-LFP amplitude across the firing-rate groups. Top: comparison of the normalized STA-LFP amplitude across the four firing-rate groups with LGN units that evoked four spikes at most within the 20-ms period. Middle: comparison of the normalized STA-LFP amplitude across the three firing-rate groups with LGN units that evoked three spikes at most within the 20-ms period. Bottom: comparison of the normalized STA-LFP amplitude across the two firing-rate groups with LGN units that evoked two spikes at most within the 20-ms period. The horizontal dashed lines represent the normalized amplitudes equal to one (i.e., same as the amplitude of the STA-LFP calculated with all spikes). Other conventions are as in Fig. 1.

The firing rate of spiking activity of an LGN unit within a 20-ms period was positively related to the STA-LFP amplitude. A representative example is shown in Figure 2*B*. When four spikes were emitted by the LGN unit within a 20-ms period, the STA-LFP amplitude was − 0.67 (a.u.) and was the largest (number of four-spikes/20 ms = 231; Fig. 2*B*, magenta line), whereas only one spike was emitted by the LGN unit within a 20-ms period, the amplitude of the STA-LFP was − 0.25 (a.u.) and was the smallest (number of one-spike/20 ms = 4978; Fig. 2*B*, red line). When two or three spikes were emitted by the LGN unit within a 20-ms period, the STA-LFP had intermediate amplitudes (− 0.37 (a.u.) for two-spikes/20 ms, number of two-spikes/20 ms = 1608, Fig. 2*B*, green line; − 0.51 (a.u.) for three-spikes/20 ms, number of three-spikes/20 ms = 666, Fig. 2*B*, blue line). The STA-LFP calculated with all spikes (number of spikes = 11,430) of the LGN unit (all-spike STA-LFP) had an amplitude of − 0.38 (a.u.; Fig. 2*B*, black line).

This pattern was confirmed with the LGN unit population. For population analysis, the amplitude of the STA-LFP of an LGN unit was normalized to the amplitude of the STA-LFP calculated with all spikes of the LGN unit (all-spike STA-LFP; Fig. 2*B*, black line, as an example). Note that normalization of the negative amplitude of STA-LFP with the negative amplitude of all-spike STA-LFP resulted in a positive value. The normalized STA-LFP amplitude was the largest (median = 1.61; number of units = 615; Fig. 2*C*, top) when LGN units emitted four spikes within a 20-ms period (four-spikes/20 ms). By contrast, the normalized STA-LFP amplitude was the smallest (median = 0.86) when the LGN units emitted only one spike within a 20-ms period (one-spike/20 ms). The normalized STA-LFP amplitude differed across the four firing-rate groups (*p* = 4.22×10^− 65^, χ² = 302, number of units = 615, Friedman’s test) and increased with the number of spikes within a 20-ms period. The normalized STA-LFP amplitude differed across the firing-rate groups for units that showed three spikes at most within a 20-ms period (*p* = 1.97×10^− 76^, χ² = 349, number of units = 1467; Fig. 2*C*, middle) and for units that showed two spikes at most within a 20-ms period (*p* = 2.96×10^− 30^, χ² = 131, number of units = 859; Fig. 2*C*, bottom). The normalized STA-LFP amplitude was the largest when the LGN units emitted the maximum number of spikes within a 20-ms period. A similar tendency was observed for spontaneous activity. These findings suggest that the firing rate of spiking activity of a single thalamic neuron within a short period was positively related to the degree of cortical activation.

### Relationship between the intervals of LGN spikes and the amplitude of cortical LFP

The relationship between the inter-spike interval (ISI) of a single LGN neuron and cortical LFP amplitude was examined. On the one hand, because the instantaneous firing rate is negatively related to ISI, a short interval between triggering and following spikes of an LGN neuron is expected to be associated with large amplitude cortical LFP. On the other hand, because the inhibitory current induced by FFI was decreased and the depression of excitatory synapses was alleviated during a long silent period of spiking activity, a long interval between the preceding and triggering spikes of an LGN neuron is expected to be associated with large amplitude STA-LFP. Each LGN spike was classified into one of five groups according to the ISIs (i.e., <20 ms, 20–100 ms, 100–200 ms, 200–500 ms, and ≥500 ms) between triggering and following spikes (following ISI; Fig 3*A*, left) or ISIs between preceding and triggering spikes (preceding ISI; Fig 3*B*, left). Although the above analysis of firing rate was examined over a 20 ms-period, the analyses with ISI were performed using a longer time scale, because inhibitory current and synaptic depression have a relatively longer time scale. LGN units evoking significant modulation in STA-LFP and emitting at least 50 triggering spikes for each of the five ISI groups were used for the analysis (number of units = 2680). The STA-LFP amplitude was normalized to the amplitude of the all-spike STA-LFP.

**Figure 3.**
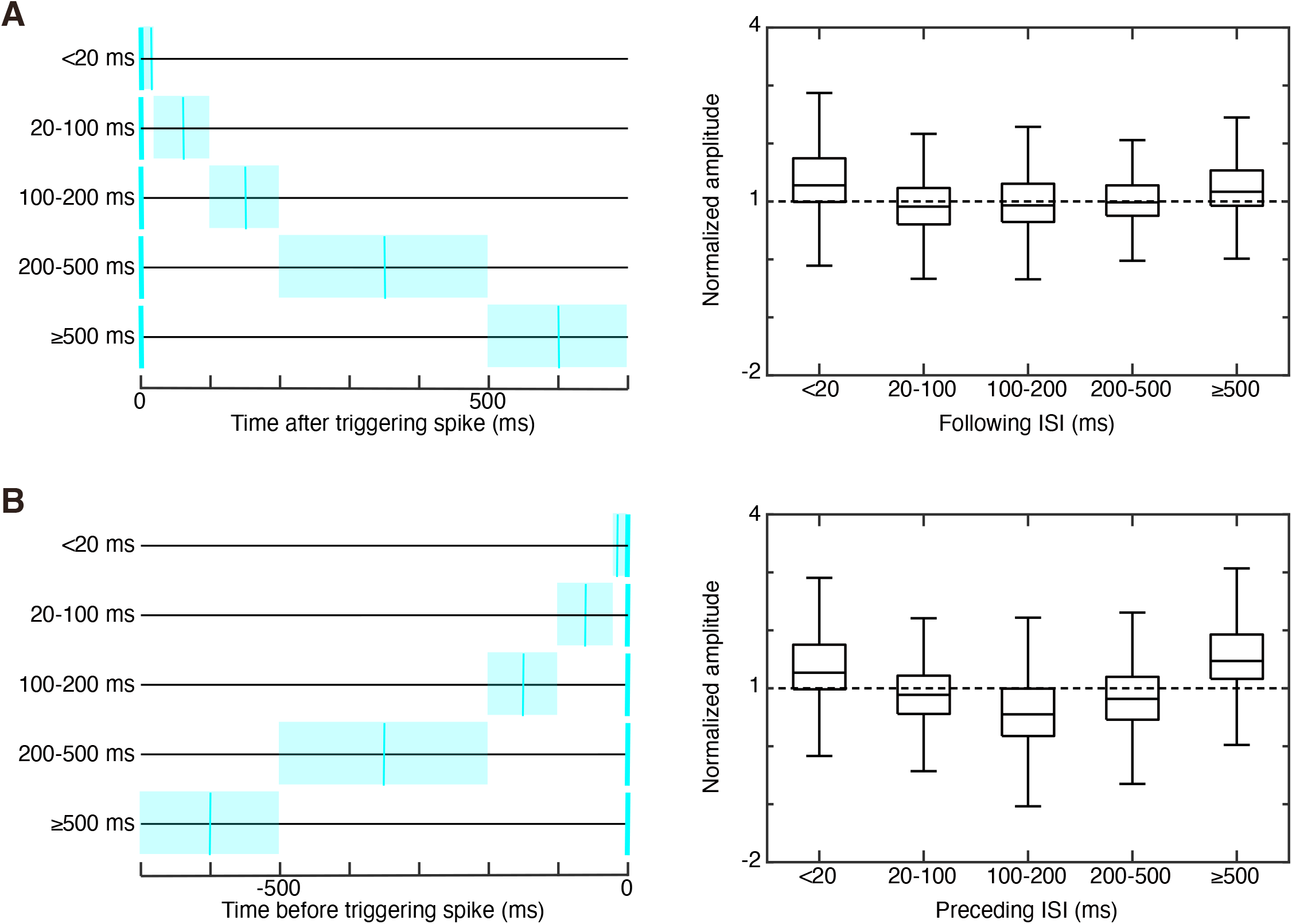
Relationship between the inter-spike interval (ISI) of spiking activity of a lateral geniculate nucleus (LGN) neuron and the amplitude of cortical local field potential (LFP). ***A***, Left: schematic depiction of intervals between triggering and following LGN spikes (following ISI, <20, 20–100, 100–200, 200–500, and ≥500 ms). ISI time windows are represented by pale cyan strips. The thick cyan vertical lines at time zero represent the timing of the triggering spike and the thin cyan vertical lines represent an example of timing of a following spike. Right: relationship between the following ISI and the normalized STA-LFP amplitude. ***B***, Left: schematic depiction of intervals between preceding and triggering LGN spikes (preceding ISI, <20, 20–100, 100–200, 200–500, and ≥500 ms). Right: relationship between preceding ISI and the normalized STA-LFP amplitude. Other conventions are as in Figs. 1 and 2.

The largest STA-LFP amplitude (median = 1.28) was induced by spikes with a following ISI of <20 ms (Fig. 3*A*, right). This finding is consistent with that obtained with firing rate during a 20-ms period (i.e., a higher firing rate during a 20-ms period of LGN spiking activity was associated with larger STA-LFP amplitude). The smallest (median = 0.91) was induced by spikes with a following ISI of 20–100 ms. The normalized STA-LFP amplitude differed across the five groups of following ISIs (*p* = 6.28×10^− 204^, χ² = 948, number of units = 2680, Friedman’s test). Results obtained with spontaneous activity were different. Although large STA-LFP amplitude (median = 1.69) was induced by spikes with a following ISI of <20 ms, the largest (median = 1.72) was induced by a following ISI of ≥500 ms. The smallest (median = 0.91) was induced by a following ISI of 100–200 ms. Although, the interval between the triggering and following spikes of an LGN unit affected the amplitudes of cortical LFP, the effect was small and inconsistent.

Next, a similar analysis was performed but the relationship between preceding ISI (interval between preceding and triggering spikes) and STA-LFP amplitude was assessed. The normalized amplitude of the STA-LFP differed across the five preceding ISI groups (*p* ≃ 0, χ² = 2865, number of units = 2680, Friedman’s test; Fig. 3*B*, right). Spikes with a long preceding ISI (≥500 ms) were associated with the largest normalized STA-LFP amplitude (median = 1.47). Spikes with a moderate preceding ISI (100–200 ms) were associated with the smallest normalized STA-LFP amplitude (median = 0.55; Fig. 3*B*). Similar results were obtained with spontaneous activity. These results suggest that the interval between the preceding and triggering spikes of an LGN unit have an impact on the amplitudes of cortical LFP.

LFP amplitude was more susceptible to the preceding ISIs than to the following ISIs. The range of normalized STA-LFP amplitudes across the ISI groups was quantified by calculating the difference between the maximum and minimum amplitudes of STA-LFP among the ISI groups. The range of the preceding ISI groups (1.69, median) was larger than that of the following ISI groups (1.31, median; effect size = 0.31, *p* = 2.72×10^− 115^, number of units = 2680, signed-rank test), suggesting that the interval between preceding and triggering spikes has a larger effect on the degree of cortical activation than the interval between triggering and following spikes.

### The temporal patterns of LGN spikes are related to the amplitude of cortical LFP

Because both preceding ISI and following ISI related to the STA-LFP amplitude, the relationship between combinations of preceding and following ISIs *and* the STA-LFP amplitude was investigated. Triggering spikes of an LGN neuron were classified into 16 groups based on combinations of preceding ISI (<20, 20–100, 100–500, and ≥500 ms) and following ISI (<20, 20–100, 100–500, and ≥500 ms). Three examples of ISI combinations are shown in Fig. 4*A*. LGN units evoking a significant modulation in the STA-LFP and emitting at least 50 triggering spikes for each of the 16 ISI combinations were analyzed (number of units = 1216).

**Figure 4.**
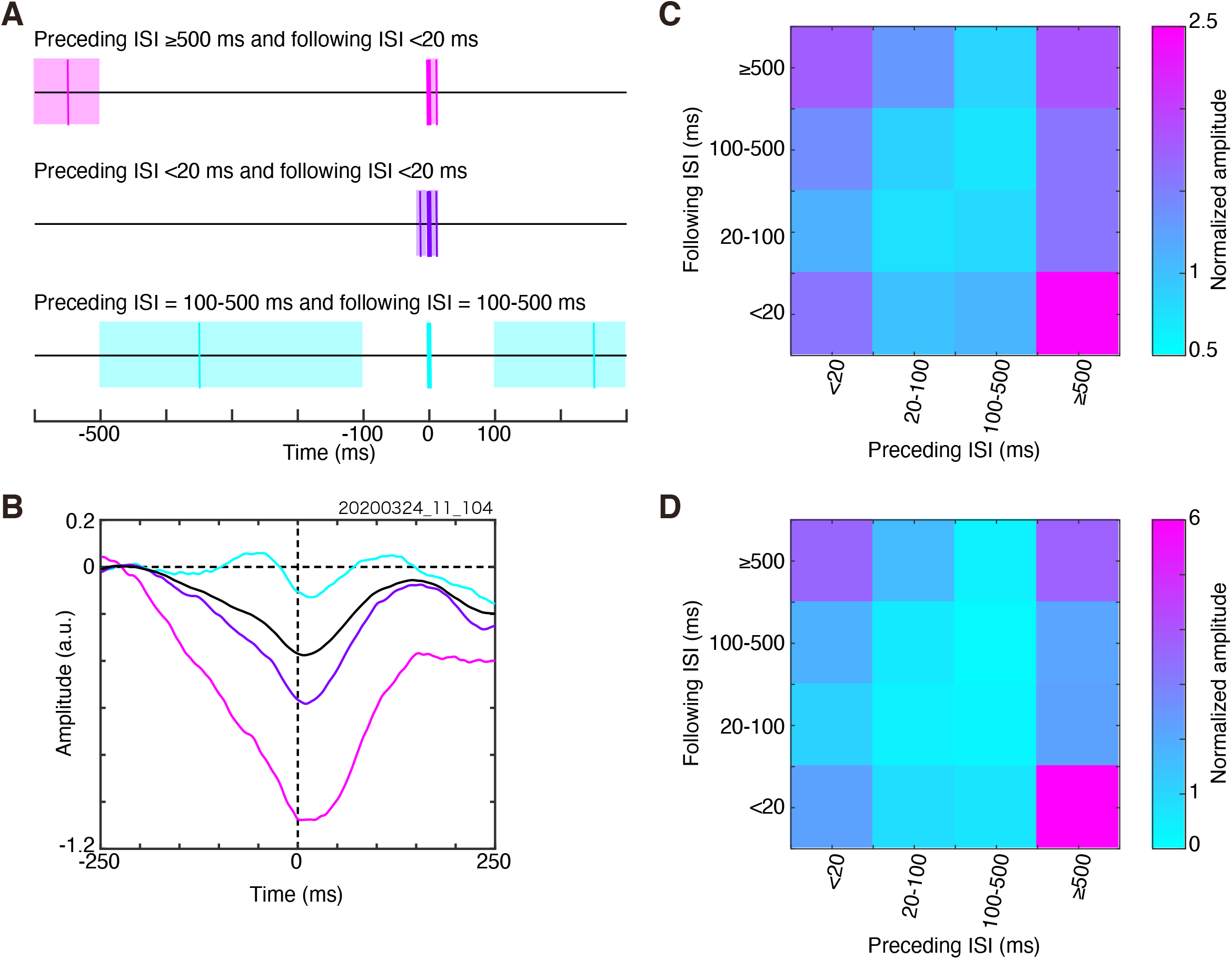
Relationship between the temporal pattern appearing in three consecutive spikes of a lateral geniculate nucleus (LGN) neuron and the amplitude of cortical local field potential (LFP). ***A***, Schematic depiction of three examples of a temporal pattern appearing in three consecutive spikes. Top (magenta): a long preceding inter-spike interval (ISI) (≥500 ms) with a short following ISI (<20 ms). Middle: (purple), a short preceding ISI (<20 ms) with a short following ISI (<20 ms). Bottom (cyan): a modest preceding ISI (100–500 ms) with a modest following ISI (100–500 ms). ***B***, An example of spike-triggered average (STA)-LFPs calculated for the three representative temporal patterns (magenta, a long preceding ISI (≥500 ms) and a short following ISI (<20 ms); purple, a short preceding ISI (<20 ms) and a short following ISI (<20 ms); cyan, a modest preceding ISI (100–500 ms) and a modest following ISI (100–500 ms)). The STA-LFP calculated with all spikes of the LGN unit (all-spike STA-LFP) was also plotted (black). This LGN unit is the same as that in Fig. 2B. ***C***, The median of the normalized STA-LFP amplitude for the 16 ISI combinations calculated using neural activity recorded during visual stimulation. The preceding ISI was plotted on the horizontal axis, the following ISI was plotted on the vertical axis, and the median of the normalized LFP amplitude was color-coded. ***D***, The median of the normalized STA-LFP amplitude for the 16 ISI combinations calculated using spontaneous activity. Other conventions are as in Figs. 1 and 3.

The normalized STA-LFP amplitude differed among the 16 ISI combinations. A representative example is shown in Figure 4*B*. When a triggering LGN spike was preceded by another spike with a long interval (≥500 ms, preceding ISI) and followed by yet another spike with a short interval (<20 ms, following ISI), the amplitude of the STA-LFP was the largest (− 1.08 (a.u.); number of spikes = 442; Fig. 4*B*, magenta line). The combination of a short preceding ISI (<20 ms) and short following ISI (<20 ms) induced STA-LFP with a moderate amplitude of − 0.58 (a.u.; number of spikes = 1667; Fig. 4*B*, purple line). A combination of moderate ISIs (preceding ISI, 100–500 ms; following ISI, 100–500 ms) was associated with the smallest STA-LFP amplitude (− 0.13 (a.u.); number of spikes = 916; Fig. 4*B*, cyan line). The STA-LFP calculated with all spikes (number of spikes = 11,430) of the LGN unit (all-spike STA-LFP) had an amplitude of − 0.38 (Fig. 4*B*, black line).

Dependence of the STA-LFP amplitude on ISI combinations was confirmed with the LGN unit population. When a triggering LGN spike was preceded by another spike with a long interval (≥500 ms, preceding ISI) and followed by yet another spike with a short interval (<20 ms, following ISI), the largest normalized STA-LFP amplitude (STA-LFP amplitude normalized to the amplitude of the all-spike STA-LFP) was observed (median = 2.46; Fig. 4*C*). A combination of moderate ISIs (preceding ISI, 100–500 ms; following ISI, 100–500 ms) resulted in the smallest normalized STA-LFP amplitude (median = 0.71). The normalized STA-LFP amplitude differed across the 16 ISI combinations (*p* ≃ 0, χ² = 4434, number of units = 1216, Friedman’s test).

Although the effect of following ISI on the amplitude of STA-LFP was weak in the previous analysis (see Relationship between the intervals of LGN spikes and the amplitude of cortical LFP), the effect of following ISI becomes obvious if analyzed in combination with the preceding ISI. For example, the largest normalized STA-LFP amplitude (median = 2.46) was observed with the long preceding ISI (≥500 ms) and the short following ISI (<20 ms), while much smaller normalized STA-LFP amplitude (median = 1.57) was observed with the same long preceding ISI (≥500 ms) but with a moderate following ISI (100–500 ms). It suggests that both preceding ISI and following ISI related to the degree of cortical activation.

If the firing rate during a short period of spiking activity of an LGN neuron is the only determinant of cortical LFP amplitude, the combination of short preceding and following ISIs is expected to result in the largest STA-LFP amplitude. Similarly, the combination of long preceding and following ISIs is likely to result in the smallest STA-LFP amplitude. However, the normalized STA-LFP amplitude induced by the combination of short preceding and following ISIs (median = 1.57) was not the largest, and the normalized STA-LFP amplitude induced by the combination of long preceding and following ISIs (median = 1.83) was not the smallest. These results suggest that the firing rate during a short period of spiking activity of an LGN neuron is not the only determinant of cortical LFP amplitude, but suggests the importance of temporal patterns appearing in three consecutive spikes (i.e., a combination of preceding ISI and following ISI) on the STA-LFP amplitude.

The above results were obtained from neural activity recorded during visual stimulation. The same analyses were performed on neural data obtained without visual stimuli (i.e., spontaneous activity), with qualitatively similar results. The normalized STA-LFP amplitude differed across the 16 ISI combinations (*p* = 7.61×10^− 259^, χ² = 1257, number of units = 166, Friedman’s test; Fig. 4*D*). The combination of a long preceding ISI (≥500 ms) and a short following ISI (<20 ms) was associated with the largest normalized STA-LFP amplitude (median = 5.98). The combination of moderate ISIs (100–500 ms, preceding ISI; 100–500 ms, following ISI) was associated with the smallest normalized STA-LFP amplitude (median = 0.17). Thus, the relationship between temporal patterns of thalamic spiking activity and cortical LFP amplitude was not limited to neural activity during sensory stimulation, but was also observed during spontaneous neural activity.

To examine whether STA-LFP associated with a specific temporal pattern could be explained by the linear summation of single-spike triggered STA-LFP, STA-LFP was reconstructed by the summation of single spike triggered STA-LFP and compared with the observed STA-LFP. This analysis focused on two ISI combinations: i) a long preceding ISI (≥500 ms) and short following ISI (<20 ms); and ii) a short preceding ISI (<20 ms) and a short following ISI (<20 ms). The former induced STA-LFP with the largest amplitude whereas the latter induced STA-LFP with a moderate amplitude, although the latter was accompanied by the highest firing rate. The analysis was performed with raw STA-LFP, because the same PSTH-predictor was subtracted from the original and reconstructed STA-LFPs.

If the STA-LFP was triggered by spikes with a long preceding ISI (≥500 ms) and a short following ISI (<20 ms), the amplitude of the observed STA-LFP was slightly larger than that of the reconstructed STA-LFP. A representative example is shown in Figure 5*A*. The amplitude of the observed STA-LFP was − 1.06 (Fig. 5*A*, red line) and that of the reconstructed STA-LFP was − 0.73 (Fig. 5*A*, blue line). This tendency was confirmed with the neuron population. The median normalized amplitude of the observed STA-LFP of the neuron population was 2.92 (median), which was slightly larger than that of the reconstructed STA-LFP (2.29, median; effect size = 0.16, *p* = 9.49×10^− 16^, number of units = 1216, signed-rank test; Fig. 5*B*).

**Figure 5.**
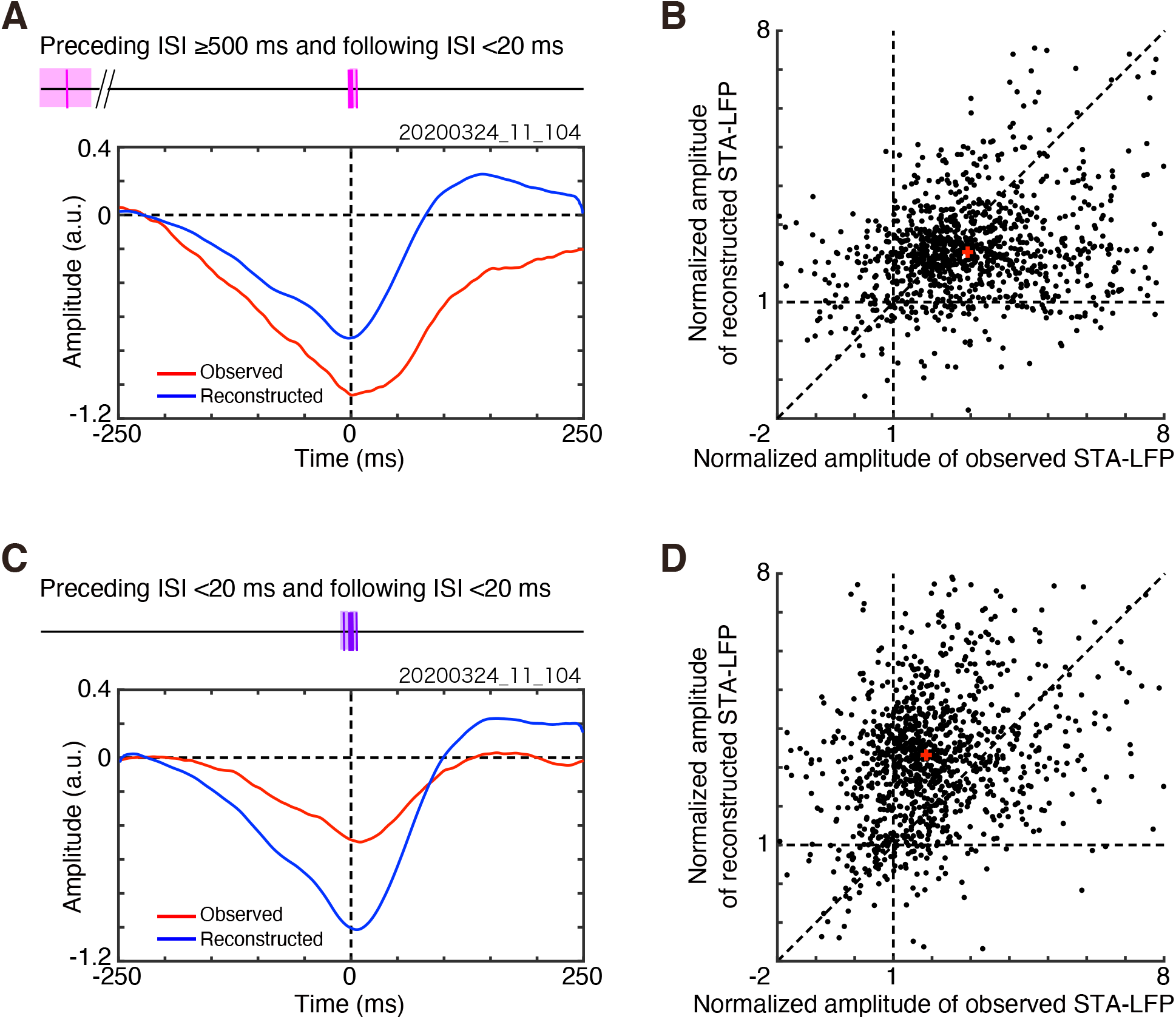
Comparison of the observed spike-triggered average (STA) of cortical local field potential (LFP) and reconstructed STA-LFP. ***A***, An example of the observed STA-LFP (red line) and reconstructed STA-LFP (blue line) for a temporal pattern with a long preceding ISI (≥500 ms) and a short following ISI (<20 ms). This LGN unit is the same as that in Fig. 4*B*. ***B***, Comparison between the normalized amplitude of the observed STA-LFP and that of the reconstructed STA-LFP for a temporal pattern with a long preceding ISI (≥500 ms) and a short following ISI (<20 ms). Each dot represents a single LGN unit. The horizontal and vertical dashed lines represent the normalized amplitudes equal to one (i.e., same as the amplitude of the STA-LFP calculated with all the spikes). The diagonal dashed line represents equality. The red cross represents the medians. ***C***, An example of the observed STA-LFP (red line) and reconstructed STA-LFP (blue line) for a temporal pattern with a short preceding ISI (<20 ms) and a short following ISI (<20 ms). This LGN unit is the same as that in A. ***D***, Comparison between the normalized amplitude of the observed STA-LFP and the reconstructed STA-LFP for a temporal pattern with a short preceding ISI (<20 ms) and a short following ISI (<20 ms). Other conventions are as in Fig. 1.

Contrary to the above results, the amplitude of the observed STA-LFP was smaller than that of the reconstructed STA-LFP if the STA-LFP was triggered by spikes with a short preceding ISI (<20 ms) and a short following ISI (<20 ms). A representative example is shown in Figure 5*C*. The amplitude of the observed STA-LFP of the LGN unit was − 0.50 (Fig. 5*C*, red line) and that of the reconstructed STA-LFP was − 1.01 (Fig. 5*C*, blue line). The median normalized amplitude of the observed STA-LFP of the neuron population was 1.85 (median) and was smaller than that of the reconstructed STA-LFP (3.32, median; effect size = 0.40, *p* = 1.70×10^− 86^, number of units = 1216, signed-rank test; Fig. 5*D*). These results suggested that the amplitude of the STA-LFP associated with spikes with a long preceding ISI and a short following ISI could be explained by the linear or supralinear summation of single-spike triggered STA-LFP, whereas that associated with a short preceding ISI and a short following ISI could be explained by the sublinear summation of single-spike triggered STA-LFP.

The amplitude of STA-LFP related to the firing rate during a 20-ms period and the largest STA-LFP amplitude were observed when triggered by the first spike of four spikes within a 20-ms period (Fig. 2). In addition, the amplitude of STA-LFP related to the temporal pattern of the spiking activity of an LGN unit and the largest STA-LFP amplitude were observed when triggered by spikes with a long preceding ISI (≥500 ms) and a short following ISI (<20 ms) (Fig. 4). The normalized amplitude of STA-LFP triggered by two types of spikes (the first spike of four spikes in a 20-ms window and the spike with a long preceding ISI (≥500 ms) and a short following ISI (<20 ms)) was directly compared. When the unit in Figure 2B emitted four spikes within a 20-ms period the STA-LFP amplitude was − 0.67 (a.u.) and when the same unit emitted spikes with a long preceding ISI (≥500 ms) and a short following ISI (<20 ms), the amplitude of the STA-LFP was − 1.08 (a.u.; Fig. 4*B*). This comparison suggested the temporal pattern of spiking activity had a much larger effect on the amplitude of cortical LFP than the firing rate within a short period. This pattern was confirmed with the neuron population. The normalized amplitude of the STA-LFP of the latter (2.18, median; preceding ISI ≥500 ms and following ISI <20 ms) was larger than that of the former (1.61, median; four spikes/20 ms; effect size = 0.28, *p* = 1.20×10^− 13^, number of units = 361, signed-rank test; Fig. 6). A temporal pattern appearing in three consecutive spikes had a much larger effect on the amplitude of the cortical LFP than the firing rate within a short period.

**Figure 6.**
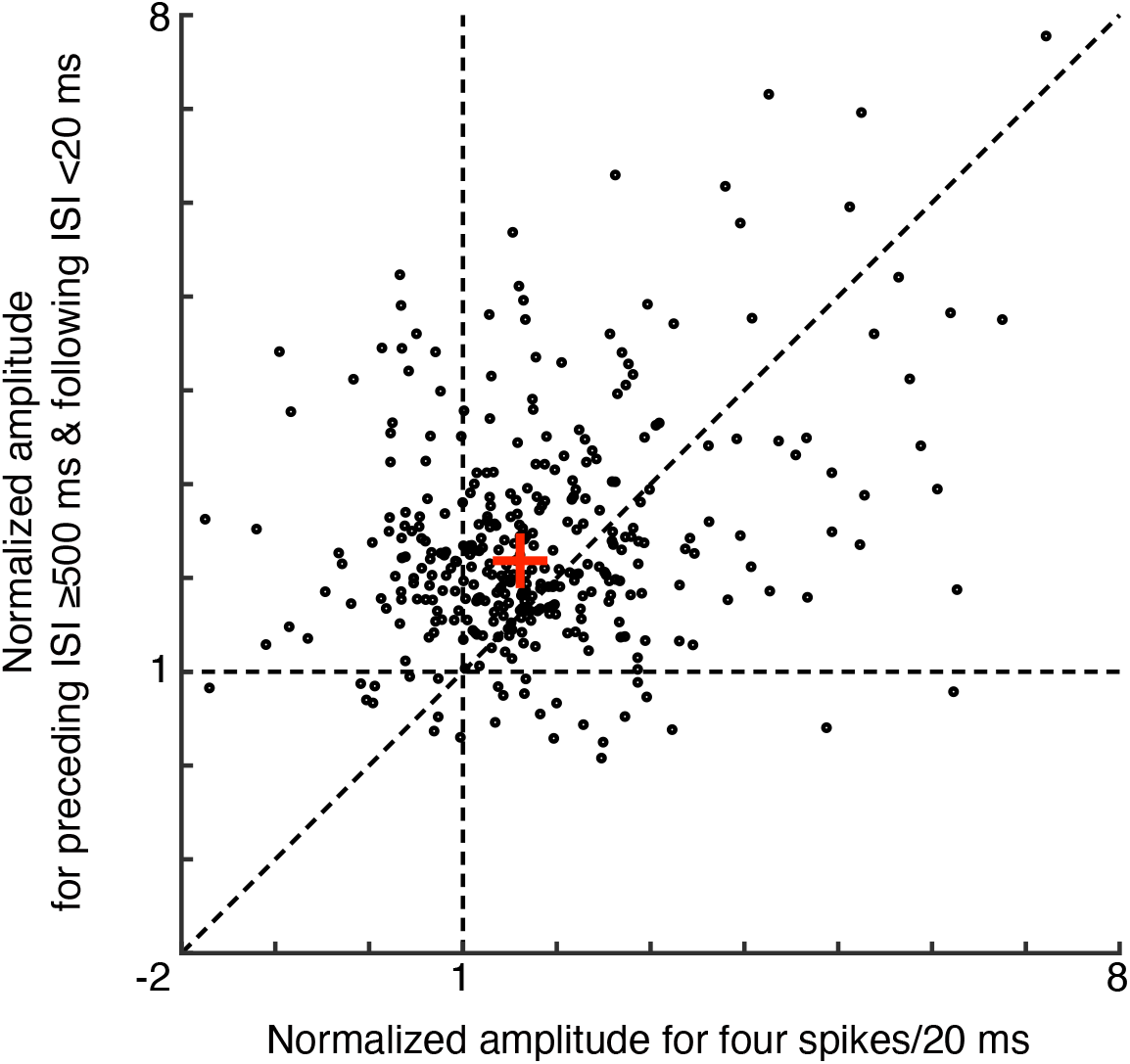
Comparison between the normalized amplitude of the spike-triggered average of local field potential induced by four spikes within a 20-ms period and that induced by a temporal pattern of spikes with a long preceding inter-spike interval (≥500 ms) and a short following inter-spike interval (<20 ms). Other conventions are as in Fig. 5.

### Spiking activity of cortical neurons and the temporal patterns of LGN spikes

To examine whether cortical spiking activity is also related to temporal pattern of spiking activity of LGN neurons, the relationship between the temporal patterns of LGN spiking activity and occurrence of cortical spiking activity was analyzed using monosynaptically connected LGN-VC unit pairs. Monosynaptically connected LGN-VC unit pairs were detected with cross-correlation analysis of their spike trains. Cross-correlation analysis was performed with LGN-VC unit pairs each having ≥5000 spikes for reliable calculation of CCG (n = 86,292 pairs). The criteria for monosynaptic excitatory connection was short latency (1–4 ms; Tanaka, 1983; Hata et al., 1990; Usrey et al., 2000; Tamura et al., 2004) significant peak (*P* < 0.0001, binomial test) in CCGs (Fig. 7*A*). Among the 86,292 LGN-VC unit pairs, 174 pairs showed the sign of monosynaptic connections.

**Figure 7.**
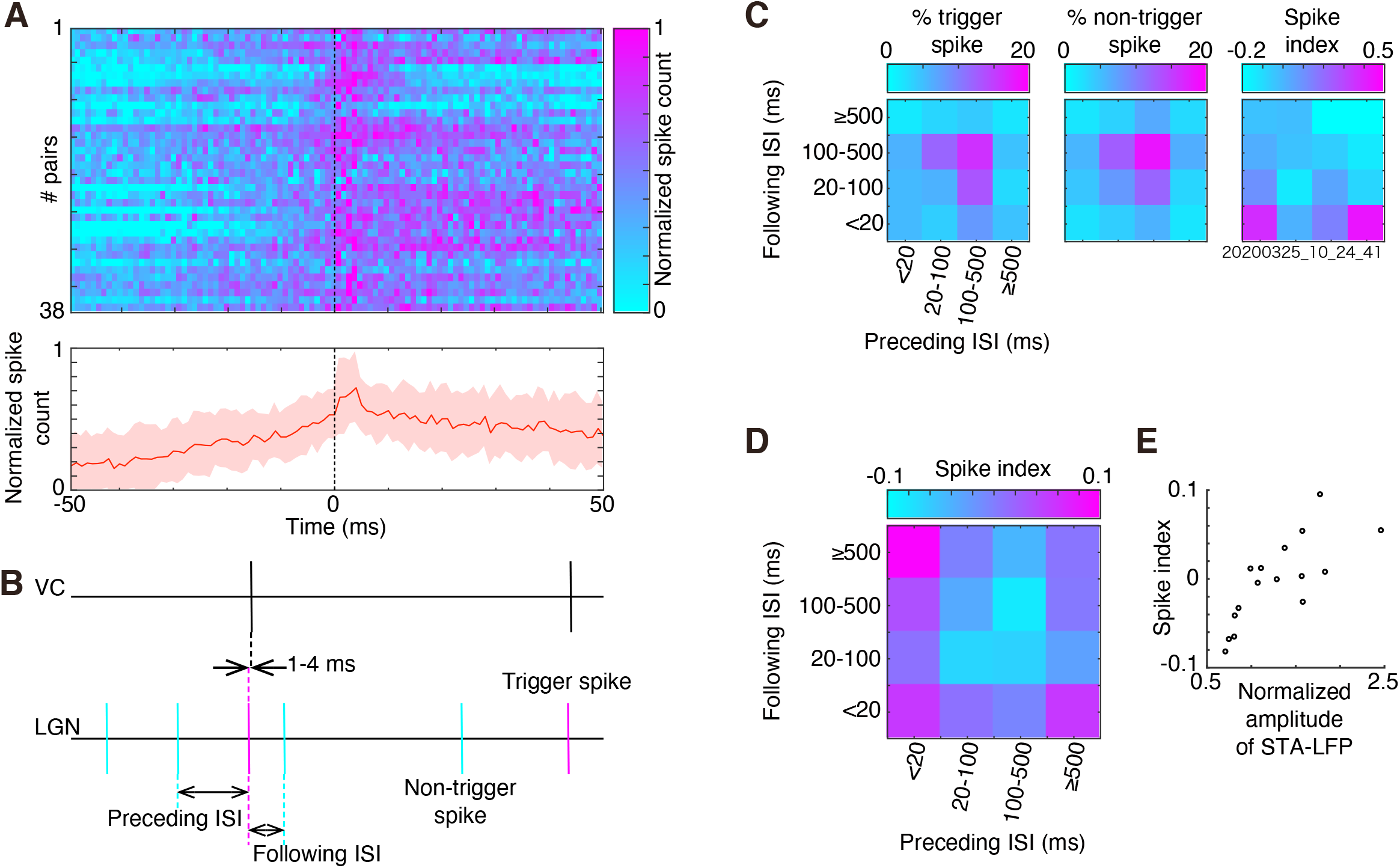
Relationship between spiking activity of a visual cortex (VC) unit and the temporal pattern of spiking activity of a lateral geniculate nucleus (LGN) unit. ***A***, Cross-correlograms (CCGs) of 38 LGN-VC unit pairs with short latency delayed peak, which suggests the monosynaptic excitatory connections from an LGN unit to a VC unit. Spike timing of a VC unit were plotted relative to spike timing of an LGN unit. Spike counts were normalized with the peak count. Top: color-coded CCGs for the 38 LGN-VC unit pairs. Each row corresponds to the CCG of a single LGN-VC unit pair. CCGs were displayed in no particular order. The normalized spike counts were color-coded. Bottom: the average of peak-normalized CCGs across the 38 pairs. Shading represents standard deviation. The vertical dashed line at the 0 ms represents the timing of LGN spikes. ***B***, Schematic depiction of classification of spikes of an LGN unit. A spike train of a VC unit (top) and an LGN unit (bottom). Each LGN spike was classified into triggering spike (magenta; an LGN spike followed by a VC spike with short latency of 1–4 ms) and non-triggering spike (cyan). The LGN spikes were further classified into 16 groups based on combination of preceding ISI (<20, 20–100, 100–500, and ≥500 ms) and following ISI (<20, 20–100, 100–500, and ≥500 ms). ***C***, An example of spike count proportion calculated with trigger spike (left) and non-trigger spike (middle) and the spikes index (right) for the 16 ISI combinations. Proportions of spike count (left and middle) and the spike index (right) were color-coded. ***D***, The median of the spike index across 38 LGN-VC unit pairs for the 16 ISI combinations. The median of the spike index was color-coded. ***E***, Relationship between the median of the spike index and the median of the normalized STA-LFP amplitude. Each dot represents a single ISI combination. Other conventions are as in Fig. 4.

Even for the monosynaptically connected LGN-VC unit pairs, not all the LGN spikes were followed by a VC spike with short latency. LGN spikes that were followed by a VC spike were designated as trigger spikes and other LGN spikes were designated as non-trigger spikes (Fig. 7B). To examine whether the trigger LGN spikes are associated with a specific combination of preceding and following ISIs, temporal patterns of three consecutive spikes around trigger spikes (preceding, triggering and following spikes) were compared with those around non-trigger spikes (preceding, non-triggering and following spikes). For this purpose, LGN spikes were further classified into 16 groups based on combinations of preceding ISI (<20, 20–100, 100–500, and ≥500 ms) and following ISI (<20, 20–100, 100–500, and ≥500 ms) (Fig. 7*B*). For each ISI combination, a spike index was calculated.

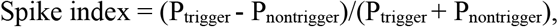

P_trigger_ is the proportion of spikes of a combination of preceding and following ISIs for trigger LGN spikes (spike count for the ISI combination divided by the number of trigger spikes), and P_nontrigger_ is that for non-trigger LGN spikes (spike count for the ISI combination divided by the number of non-trigger spikes). Positive spike index means that the ISI combination appeared more frequently around trigger spikes than non-trigger spikes. Among the monosynaptically connected LGN-VC unit pairs, LGN units emitting ≥300 trigger and non-trigger spikes were analyzed (number of LGN-VC unit pairs, 38; the number of trigger LGN spikes, 300–752; the number of non-trigger LGN spikes, 4503–25,101).

The combination of a long preceding ISI (≥500 ms) and a short following ISI (<20 ms) had a positive spike index. For example, a combination of moderate ISIs (preceding ISI, 100–500 ms; following ISI, 100–500 ms) observed most frequently (16.6 percent and 18.2 percent of trigger and non-trigger LGN spikes, respectively) in an LGN-VC unit pair (the number of trigger spikes, 386; the number of non-trigger spikes, 7837; Fig. 7*C*), and the spike index was negative (−0.06). The combination of a long preceding ISI (≥500 ms) and a short following ISI (<20 ms) was observed in 4.9 percent and 1.9 percent of trigger and non-trigger spikes of the LGN unit, respectively, and results in the spike index of 0.43, meaning that the combination appeared more frequently around trigger spikes than non-trigger spikes of the LGN unit. The median of the spike index across LGN-VC unit pairs (n = 38) for the combination of a long preceding ISI (≥500 ms) and a short following ISI (<20 ms) was 0.054 (Fig. 7*D*). The spike index differed across the 16 ISI combinations (*p* = 1.55×10^− 7^, χ ² = 61.23, number of LGN-VC unit pairs = 38, Friedman’ s test). Furthermore, the median of spike index positively correlated with the median of normalized amplitudes of STA-LFP calculated for combinations of preceding and following ISIs (*r* = 0.80, *p* = 0.00034, number of ISI combinations = 16, Spearman’ s correlation coefficient; Fig. 7*E*; see Fig. 4*C*). The results suggest that cortical spiking activity is also related to the temporal pattern of three consecutive LGN spikes in a similar manner to cortical LFP.

## Discussion

The main finding of the present study is that a specific temporal pattern appearing in three consecutive spikes of an LGN neuron induced larger LFP modulation in VC than high-frequency spiking activity during a short period. The findings indicate the importance of the temporal pattern of spiking activity of a single thalamic neuron on cortical activation.

### Technical considerations

In the present study, visual stimuli were presented to enhance spiking activity of LGN neurons and obtain reliable STA-LFP from a limited recording duration. Because presentation of visual stimuli modulates the firing rates of spiking activity of an LGN unit and cortical LFP almost simultaneously, the raw STA-LFP reflects the correlation related to the stimulus-locked modulation (stimulus coordination) *and* the correlation related to the interaction between the LGN unit and cortical LFP (neural correlation). The effect of stimulus coordination was estimated by calculating the PSTH-predicted STA-LFP and removed by subtracting the PSTH-predicted STA-LFP from the raw STA-LFP. Spontaneous neural activity was also analyzed and demonstrated similar results. Therefore, any effect of visual stimulation on the present findings was small or negligible.

Animals were anesthetized with urethane and the head of the animal was fixed to achieve stable recordings with silicon microelectrodes. These procedures may have affected the interpretation of the present findings. However, no difference in thalamocortical transmission under different states of alertness (Stoelzel et al., 2009) and the response properties of neurons in the LGN and VC of anesthetized mice were previously reported to be qualitatively similar to those of awake animals (Durand et al., 2016). Although similar conclusions are likely to be obtained with awake and behaving animals, future studies in awake and behaving animals may be required.

### STA-LFP and thalamocortical neuron

In the present study, the LFP was recorded as a measure of cortical activity, and modulation in STA-LFP was observed in a wide cortical region (>1.4 mm). The results may not be consistent with the lateral extent of a single thalamocortical axon terminal (<1.0mm, Raczkowski and Fitzpatrick, 1990). However, because LFP signals represent neural activity from a few-millimeter region of the cortex (Mitzdorf, 1987; Logothetis, 2003; Kreiman et al., 2006; Jin et al., 2008; Nauhaus et al., 2009; Kajikawa and Schroeder, 2011; Buzsáki et al., 2012; Pesaran et al., 2018), signals related to spiking activity of a single LGN neuron can be recorded from widespread cortical regions.

An LGN unit that induces modulation in cortical LFP is likely to be a thalamocortical relay neuron (TC neuron). Neurons in the LGN are divided into TC neurons and local inhibitory interneurons. TC neurons form 90% of the LGN neurons in rodents (Guido, 2018), and have a slightly broader spike width and larger soma than interneurons (Williams et al., 1996). In the present study, a large proportion of LGN units evoked significant modulation in the STA-LFP. LGN units with a broader spike width and lower firing rate, which are signatures of a larger neuron in general, induced the larger modulation of the STA-LFP. These results are consistent with the notion that an LGN unit that induces a large modulation in the STA-LFP is a TC neuron.

### Firing rate of thalamic spiking activity during a short period and cortical LFP

There was positive correlation between the firing rate of spiking activity of an LGN unit during a short period and the LFP amplitude recorded from the VC. The temporal summation of the depolarizations induced by consecutive spikes of a single LGN neuron underlies this relationship. The result is consistent with a previous study reporting a potential temporal summation at thalamocortical synapses (Usrey et al., 2000), but may not be consistent with the presence of FFI and depression of excitatory thalamocortical synapses. FFI can be weakened by the repetitive activation of the thalamocortical pathway (i.e., depression of FFI; Gabernet et al., 2005), and the depression of FFI facilitates the temporal summation at the thalamocortical synapses.

Although high-frequency spiking activity during a short period of an LGN unit induced larger LFP modulation than single spiking activity, the increase in the LFP amplitude was limited. For example, one may expect that the amplitude of LFP induced by four consecutive spikes was about four times that induced by a single spike. However the amplitude of LFP induced by four consecutive spikes was less than two times that induced by a single spike, suggesting that the temporal summation is not as effective as the expectation. Therefore, it is reasonable to conclude that FFI was not fully suppressed and depression of excitatory thalamocortical synapses was not fully recovered. High frequency spiking activity of a LGN neuron induced successive depolarization in cortical neurons under influence of FFI and depression of excitatory thalamocortical synapses, and results in a weak but larger depolarization than a single LGN spike.

### Temporal pattern of thalamic spiking activity and cortical LFP

Temporal pattern appearing in three consecutive spikes of an LGN unit was related to the STA-LFP amplitude as well as to spiking activity of cortical neurons. A large modulation of the STA-LFP was observed, if a triggering LGN spike was preceded by another spike with a long interval (long preceding ISI) and was followed by yet another spike with a short interval (short following ISI). Contrary, the combination of a moderate ISIs was associated with the smallest normalized STA-LFP amplitude. Consistent results were obtained with the analysis of spiking activity of a cortical unit receiving monosynaptic inputs from an LGN unit. Thus, it is concluded that combination of preceding ISI and following ISI related to the degree of cortical activation. Consistent with the present results, it has been show that the preceding ISI of a thalamic neuron are related to the degree of cortical activation (Swadlow and Gusev, 2001; Swadlow, 2002; Swadlow et al., 2002; Stoelzel et al., 2008; Stoelzel et al., 2009).

Importance of temporal pattern appearing in three consecutive spikes of an LGN neuron on cortical activation is not simply reflecting the effect of firing rate of an LGN neuron on cortical activation. If the combination of short preceding and following ISIs induced the largest LFP modulation and the combination of long preceding and following ISIs induced the smallest LFP modulation, one may conclude that the firing rate of a thalamic neuron during a short period was the determinant of the degree of cortical activation. However, these were not the case, suggesting that the firing rate during a short period is not the only determinant of the degree of cortical activation. Furthermore, and more importantly, a temporal pattern appearing in three consecutive spikes of an LGN unit (long preceding ISI and short following ISI) induced larger cortical LFP modulation than high-frequency spiking activity during a short period, suggesting the importance of temporal pattern that is unrelated to firing rate of an LGN neuron on cortical activation.

The amplitude of the STA-LFP associated with spikes with a long preceding ISI and a short following ISI was explained by the linear or supralinear summation of single-spike triggered STA-LFP. During the long-preceding silent period of a LGN neuron, its thalamocortical excitatory synapses recovered from the depressed state and the inhibitory currents induced by FFI were decreased. Then, depolarization induced by triggering and following LGN spikes are effectively and (supra)linearly summated in cortical neurons. Contrary, the amplitude of the STA-LFP associated with spikes with a short preceding ISI and a short following ISI was explained by the sublinear summation of single-spike triggered STA-LFP. A spike induces excitatory currents as well as FFI currents in cortical neurons *and* depresses its related synapses. Then, LGN spikes following with short interval induce much smaller depolarization than the first and result in sublinear summation in cortical neurons. These results can be interpreted that preceding ISI controls the gain of depolarization induced by following spikes.

Although the present results clarify the importance of temporal pattern appearing in consecutive spikes of a single thalamic neuron on the thalamocortical transmission, the results did not deny the contribution of spatial summation at the thalamocortical transmission. Synchronous spiking activity between thalamic neurons has been reported and implicated in thalamocortical synaptic transmission (Alonso et al., 1996; Roy and Alloway 2001; Bruno and Sakmann, 2006; Ito et al., 2010). Synchronous inputs from thalamic neurons are summated spatially and induce large depolarization in cortical neurons. The present results are not incompatible with the potential involvement of synchronous spikes but suggest that synchronous spikes are more effective if they occur in a specific temporal pattern.

## Acknowledgments

I thank Dr. Fumitaka Kimura for his encouragement and for critically reading the manuscript, and J. Ludovic Croxford, PhD, from Edanz (https://jp.edanz.com/ac) for editing a draft of this manuscript. This work was supported by Grant-in-Aids for Scientific Research on Innovative Areas— “Innovative SHITSUKAN Science and Technology” (15H05921) and “Non-linear Neuro-Oscillology” (16H01612) from MEXT, Japan— and by a Grant-in-Aid for Scientific Research (C) (17K07056).

